# Mevalonate pathway activation in Ewing sarcoma reveals a 3D-specific synergy between statins and BCL-xL inhibition

**DOI:** 10.1101/2025.11.20.689456

**Authors:** Branka Radic-Sarikas, Marica Markovic, Caterina Sturtzel, Mathias Ilg, Martha Magdalena Zylka, Didier Surdez, Martin Metzelder, Martin Distel, Aleksandr Ovsianikov, Florian Halbritter, Heinrich Kovar

## Abstract

Bone sarcomas are rare and aggressive pediatric cancers with limited progress in targeted therapy development, in part due to the poor physiological relevance of conventional two-dimensional (2D) culture systems used for preclinical testing. To address this gap, we developed a standardized three-dimensional (3D) culture and drug-testing platform for Ewing sarcoma (ES) and osteosarcoma (OS) that more accurately recapitulates in vivo tumor biology. Across 3D spheroids, bioprinted constructs, and patient-derived xenograft (PDX) cultures, we observed a consistent activation and dependency on the mevalonate pathway in ES. Leveraging this platform, we identified a selective therapeutic synergy between statins, which inhibit mevalonate pathway flux, and BCL-xL inhibitors, a vulnerability that was not detectable in 2D cultures. These findings highlight the mevalonate pathway as a targetable metabolic dependency in ES and demonstrate how physiologically grounded 3D models can uncover clinically actionable treatment strategies that remain hidden in traditional 2D systems. Our findings show that 3D tumor models can expose actionable metabolic vulnerabilities obscured by traditional approaches, supporting their use in rational combination therapy discovery for aggressive pediatric sarcomas.

## Introduction

Ewing sarcoma (ES) and osteosarcoma (OS) are the most common malignant bone tumors in children and adolescents. In ES, the primary pathogenic driver is a gene rearrangement between *EWSR1* and an ETS family gene, most commonly *FLI1*, leading to the expression of an aberrant chimeric transcription factor, EWS::FLI1 that disrupts the regulation of hundreds of genes and mediates malignant transformation. Aside from the *EWSR1::ETS* translocations, only a few other recurrent somatic alterations are observed in ES^1–3^. In contrast, OS is driven by TP53 and RB1 inactivation^4^, and its genomic complexity has been attributed either to early catastrophic events followed by stable genome, or to ongoing chromothriptic processes contributing to clonal evolution and tumor heterogeneity.^5^

Bone sarcomas are highly aggressive: 20-25% of patients present with metastatic disease at diagnosis, often resistant to intensive multi-modal therapies, leading to poor survival rates of only 20-30%^6–9^. Despite the urgent need for more effective therapeutic strategies, preclinical development remains limited by the scarcity of robust genetic animal models and the limited availability of preclinical testing platforms.

The challenges in bone sarcoma research mirror broader issues in the drug discovery process, which largely relies on two-dimensional (2D) monolayer models that fail to replicate tumor gene expression and drug response^10^. Even promising compounds frequently fail in clinical trials, with overall failure rates exceeding 90%^11^, reflecting the limitations of preclinical models. Patient-derived xenografts (PDXs) closely replicate human tumor biology, but are constrained by experimental feasibility, limited sample availability, and ethical considerations.

Developing advanced three-dimensional (3D) in vitro systems provides an attractive way to culture PDXs while retaining their biological fidelity, reducing reliance on animal studies, and aligning with the principles of replacement, reduction, and refinement (the “3Rs”), which aim to minimize animal use in research while improving experimental design ^12^. These systems provide controlled platforms for mechanistic studies and drug testing, complementing in vivo research. Optimized 3D culture systems – including scaffold-based constructs, microfluidic devices, spheroids, and organoids – provide physiologically relevant alternatives. Spheroids, in particular, replicate 3D architecture, cellular heterogeneity, hypoxic gradients and chemoresistance akin to that observed in patients^13^, while supporting high-throughput drug testing, a significant advantage over many other 3D systems. However, their utility depends on tightly controlled parameters, such as culture duration and cell density^13,14^.

While spheroids provide a versatile, scaffold-free platform, scaffold-based 3D cultures rely on appropriate extracellular matrix (ECM) support to maintain structural integrity. Traditionally, Matrigel – a mouse chondrosarcoma-derived ECM – has been used for this purpose, but its variable composition can lead to inconsistent outcomes^15^. To address this limitation, recent innovations in modular synthetic hydrogels offer a more controlled and reproducible alternative. These hydrogels accurately mimic essential ECM properties, facilitating reliable stem cell expansion and organoid formation^16–18^. Scaffold-based approaches, whether prefabricated or decellularized, involve trade-offs: prefabricated constructs allow controlled manufacturing but lack native tissue structures^19,20^, while decellularized scaffolds better mimic cell-matrix interactions but pose species-specific and contamination challenges and have limited mechanical adaptability^21^. Synthetic hydrogels thus provide a flexible, reproducible platform for scaffold-based modeling, complementing scaffold-free spheroids and expanding the toolkit for precise drug evaluation.

Here, we evaluated scaffold-free and scaffold-based 3D culture approaches in cell lines and patient-derived xenografts, establishing robust and adaptable platforms for bone sarcoma modeling and drug screening. These 3D systems revealed potent, selective drug synergies in Ewing sarcoma, including co-targeting of the mevalonate pathway and BCL-xL. Importantly, the observed drug synergy was highly context dependent, emphasizing how cellular microenvironments shape drug responses^22,23^. By testing drug combinations in physiologically relevant models, we were able to identify strategies that enhance therapeutic efficacy, highlighting the value of advanced preclinical models for optimizing treatment strategies. Our findings provide new insights into the molecular vulnerabilities of bone sarcomas and inform the development of more effective combination therapies.

## Results

### Spheroid formation and transcriptomic profiling reveal tumor-like metabolic reprogramming

To lay the groundwork for a reliable 3D drug screening platform, we first assessed spheroid formation and drug response across a panel of ES and OS cell lines. Key parameters such as culture duration and cell density were systematically evaluated using the hanging droplet^24,25^ and liquid overlay methods^26^. The liquid overlay method was selected based on reproducibility and scalability and was further refined to ensure consistent spheroid formation.

To establish a robust foundation for the platform, we initially screened four ES and six OS cell lines for their ability to consistently form uniform spheroids under low-attachment conditions. From this panel, we selected two ES lines (SK-N-MC and STA-ET-1) and two OS lines (OS143B and STA-OS-5) that demonstrated the most reliable and reproducible spheroid-forming capabilities. Although other lines exhibited partial spheroid-forming capacity, we prioritized these four models for downstream optimization due to their consistent growth characteristics.

The selected ES lines offer complementary biological relevance: STA-ET-1 line, expresses wildtype TP53^27^ reflecting the broader ES patient population, while SK-N-MC is a well-established TP53-deficient model^27^ with a more aggressive phenotype. Among the OS models, OS143B, a Ki-Ras–transformed HOS cell line derivative, is highly tumorigenic and models advanced disease^28^, while STA-OS-5, with defined chromosomal gains, provides a distinct genomic context^29^ and may model a less aggressive counterpart to OS143B.

Building on the protocol by Friedrich et al.^26^, we implemented several key modifications to adapt and streamline the workflow for bone sarcoma models (see Methods). We replaced the labor-intensive acid phosphatase assay with the CellTiter-Glo 3D assay, which required careful optimization to ensure compatibility with spheroid size and media volume. We also refined the agarose preparation, eliminated autoclaving steps without compromising sterility, and limited cell passage to ≤10 to reduce variability and avoid artifacts from prolonged 2D culturing.

Importantly, we optimized conditions to consistently generate a single, uniform spheroid per well – a critical feature for reliable drug testing, as multiple spheroids per well, a common issue in 3D culture systems, can lead to variability in drug exposure and complicate data interpretation. This was achieved by fine-tuning seeding densities, surface coating parameters, and selecting cell lines with robust spheroid-forming capacity. Spheroids consistently formed within 3–4 days, enabling treatment initiation once a structurally mature and reproducible spheroid had developed. By contrast, short-term setups produce loose aggregates that lack uniformity, making drug response measurements less reliable. Spheroids reached ∼400 µm by day 4, when treatment was initiated, and untreated spheroids grew to ∼650 µm after 72h. At this stage, spheroids remain sufficiently compact for effective drug penetration, as confirmed by doxorubicin autofluorescence in central optical sections of confocal Z-stacks (**Fig. S1A**), while still maintaining minimal central necrosis^13,26^. Cryosections confirmed that the spheroids were viable and proliferative at the start of the assay (**Fig. S1B**), an observation also supported by the continued diameter increase of untreated spheroids.

Chemotherapy treatment induced notable changes in both spheroid morphology and viability, supporting the model’s relevance for recapitulating tumor responses. As illustrated for vincristine, treatment disrupted spheroid integrity and led to a reduction in size (**Fig. S1C**). We monitored STA-ET-1 spheroids over 16 days, with doxorubicin treatment initiated on day 4 post-formation (d4). A 5-fold serial dilution (20 µM to 0.09 nM) produced clear dose-dependent growth inhibition and morphological changes (**Fig. 1A**). Untreated spheroids continued to expand, while lower drug concentrations resulted in mild but progressive growth inhibition. At the highest doxorubicin concentrations (20 µM and 4 µM), spheroids paradoxically appeared larger on day 7 despite reduced viability, consistent with oncosis – a form of cell death associated with cellular swelling and osmotic dysregulation^30–32^. These concentrations were therefore excluded from IC₅₀ determination to prevent distortion of the fitted dose–response curve. By day 14, spheroids treated with these doses had fully disintegrated, confirming potent cytotoxicity. Still, caution is warranted that spheroid size alone may not always reflect viability – particularly for drugs with complex dose-dependent effects like doxorubicin – and additional parameters such as intensity differences within a spheroid that reflect apoptotic areas should be integrated. Nonetheless, A 72-hour treatment initiated on day 4 yielded consistent, concentration-dependent responses, with IC_50_ values derived from spheroid size (**Fig. 1B**) closely matching those from viability assays (**Fig. 1C**), both of which were higher than IC_50_ obtained from 2D cultures, highlighting the tendency of monolayer models to overestimate drug efficacy.

**Figure 1.**
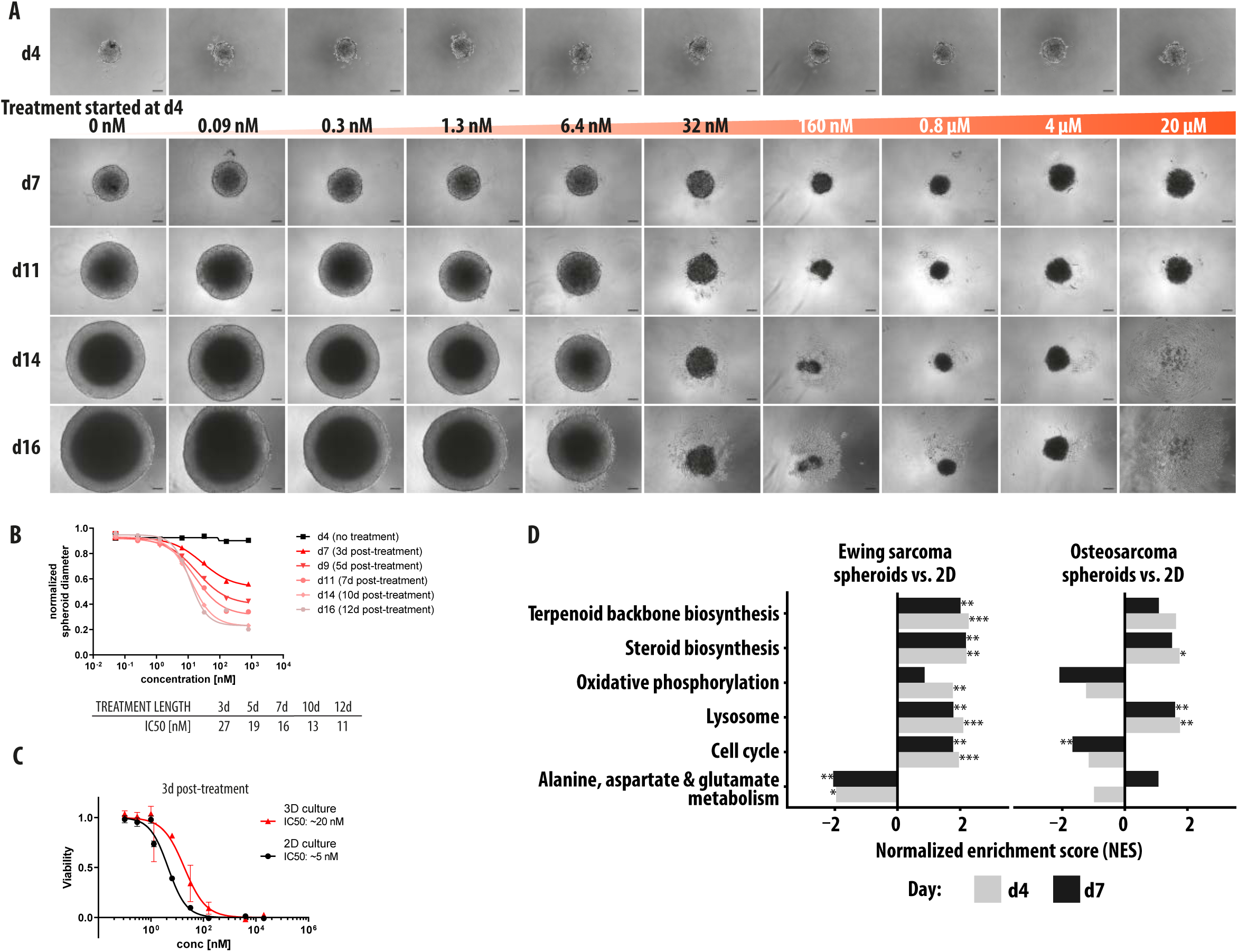
Growth dynamics, drug response, and transcriptional shifts in sarcoma spheroids. **(A)** Overview of the 16-day STA-ET-1 spheroid treatment protocol with doxorubicin, initiated on day 4 (d4). Bright-field images were acquired on days 4, 7, 11, 14, and 16, with day 7 chosen as the primary time point for downstream analyses. Notably, spheroid size decreased with increasing doxorubicin concentration across most doses, but at the two highest concentrations sizes transiently increased relative to the expected trend, likely reflecting cellular swelling (oncosis) rather than proliferation. **(B)** STA-ET-1 spheroid diameter measurements following doxorubicin treatment, obtained from bright-field images, are plotted. IC₅₀ values were derived from the corresponding dose-response curves. **(C)** Dose-response curves of doxorubicin in STA-ET-1 cells, assessed by viability assay 72 hours post-treatment, reveal a four-fold higher sensitivity in 2D cultures compared with 3D spheroids. **(D)** KEGG enrichment analysis of ES spheroids at days 4 and 7 compared to 2D cultures reveals a marked upregulation of the mevalonate pathway indicated by steroid biosynthesis and terpenoid backbone biosynthesis. Asterisks denote statistical significance based on adjusted p-values: *p* < 0.05 (\**), p < 0.01 (**), p < 0.001 (*\*\*\*).

We collected ES and OS spheroids at day 4 (d4, ∼400 µm diameter) and day 7 (d7, ∼700 µm diameter) and performed RNA-seq to compare transcriptional profiles with 2D cultures grown in identical medium. In both d4 and d7 spheroids, we found lysosomal activity was upregulated in both ES and OS spheroids (fgsea^33^, FDR-adjusted P-value <= 0.005; **Figs. 1D, S2**), reflecting adaptation to metabolic stress and tissue-like organization^34–37^. The change in environment also impacted the expression of many matrisome genes – encoding core ECM components and regulators – and secreted factors in OS spheroids, while in ES spheroids only selected ECM glycoproteins and secreted factors like *Il23A*, *BMP6*, or *TNFSF12* were upregulated (annotated in MatrisomeDB^38^; **Fig. S3)**. Intriguingly, genes related to cell cycle and oxidative phosphorylation were only upregulated in ES spheroids but downregulated in OS spheroids. Conversely, there was a downregulation of genes related to alanine, aspartate, and glutamate metabolism in ES spheroids, which was not present in OS spheroids.

Most strikingly, we observed upregulation of the mevalonate pathway in ES spheroids (“steroid biosynthesis” and “terpenoid backbone biosynthesis”; **Figs. 1D, S2**). This pathway is known to support tumor growth and progression^39–42^, including in Ewing sarcoma^43^ and its deregulation can impact lipid synthesis and cell membrane integrity, e.g., via cholesterol and isoprenoids which are key precursors for protein prenylation, cellular signaling, and metabolic adaptation. A detailed schematic of the pathway (**Fig. S4A**) highlights the differential expression of key enzymes in ES spheroids relative to 2D cultures, showing widespread transcriptional deregulation with a predominance of upregulated nodes. In contrast, OS models displayed a more restricted and distinct pattern, with fewer pathway components affected (**Figs. S2, S4B)**. Notably, OS 3D cultures showed marked upregulation CYP27B1, indicating activation of vitamin D metabolism, whereas ES spheroids showed coordinated induction of multiple cholesterol biosynthesis and handling genes, reflecting distinct, lineage-specific metabolic adaptations (**Fig. S4B**).

### Bioengineered 3D culture enables long-term culture and drug sensitivity profiling of ES models

To expand our 3D platform beyond spheroid (scaffold-free) models and enable more controlled long-term cultures, we adapted hydrogel-based systems for ES cells. Instead of seeding cells onto prefabricated constructs, we embedded them within cross-linked hydrophilic polymer networks^44,45^ (**Fig. 2A**). This method allowed homogeneous cell distribution and high initial cell density. We first optimized the composition of the bioink – a liquid mixture comprising cells, hydrogels and nutrients – by using ES cells lines with distinct molecular profiles and growth requirements (A673, TC-71 and STA-ET-1). We systematically varied technical parameters, including the type and concentration of photocrosslinkable gelatin derivatives, hydrogel concentrations, degrees of functionalization, and the types and concentrations of photoinitiators. Compatibility of different bioink formulations was evaluated using laser scanning microscopy (LSM) combined with live/dead staining (CalceinAM/PI), allowing quantitative assessment of cell viability and morphology within the 3D constructs. The best results were achieved with embedding in gelatin-norbornene (gelNB92, 7.5%) hydrogel with equimolar concentrations of crosslinker dithiothreitol (DTT) and 0.1 mM Li-TPO-L as a photoinitiator in case of one photon polymerization (UV), which enabled long-term culture of ES cells for up to 14 days (**Fig. 2B**). This optimized bioink composition was also compatible with two-photon polymerization (2PP)^46^, with 2PP requiring a different photoinitiator (DAS^47^) and allowing fabrication of constructs with higher spatial resolution and defined geometries. (**Fig. 2C**).

**Figure 2.**
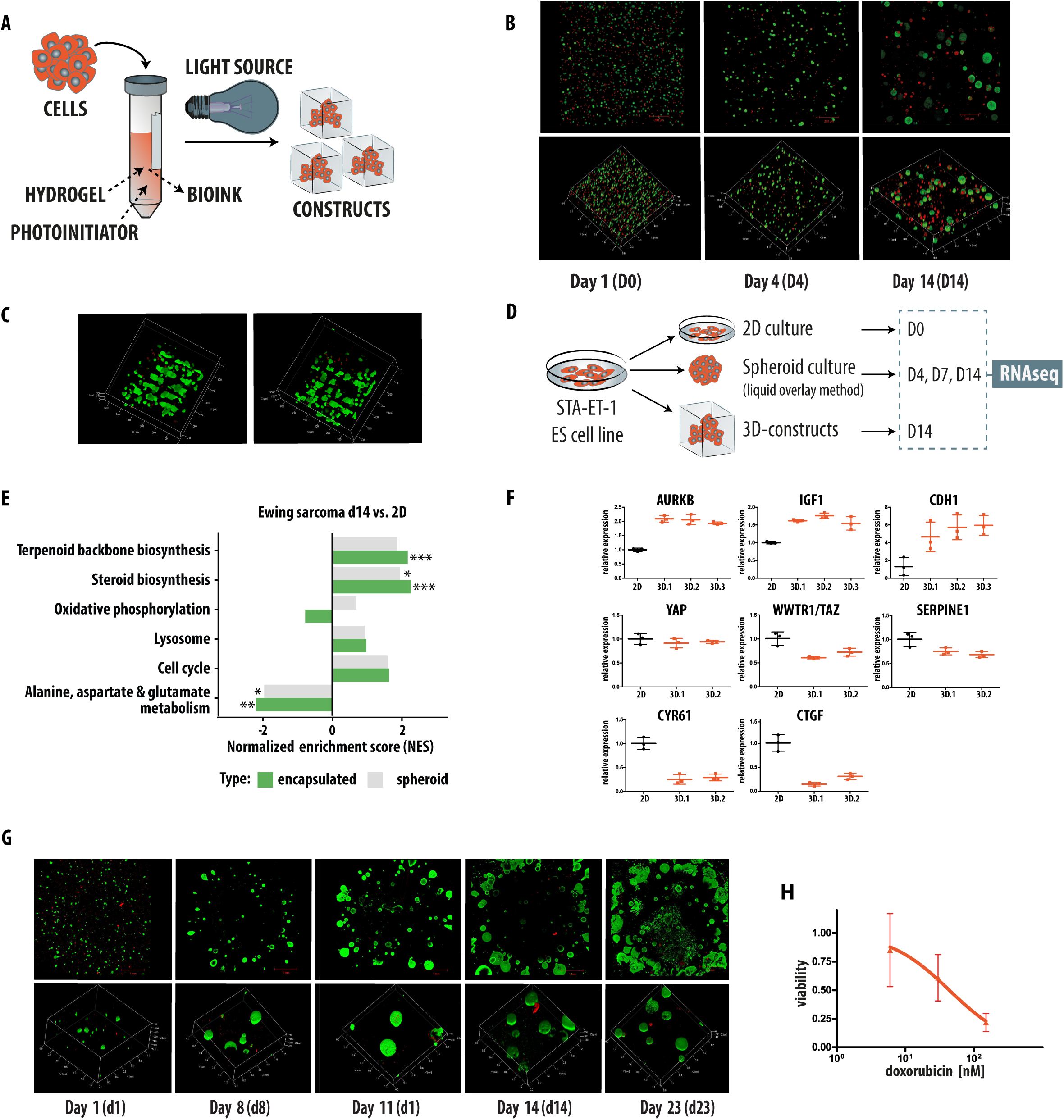
Hydrogel-based 3D culture system supports long-term Ewing sarcoma growth and drug testing. **(A)** Schematic of the embedding process for creating 3D-constructs (modified from Markovic et al.^20^): ES cells are embedded in photocrosslinkable hydrogels and polymerized to form cell-laden constructs. **(B)** STA-ET-1 cells embedded in gelNB92, 7.5% hydrogel with equimolar concentrations of crosslinker dithiothreitol (DTT) and 0.1 mM Li-TPO-L as a photoinitiator and cultured for 14 days as 3D-constructs. Cell viability was assessed using LIVE/DEAD staining, with green fluorescence indicating live cells, and red fluorescence indicating dead cells. **(C)** ES TC71 cells 48h post-embedding, a critical early period for assessing viability and structural integrity, in gelNB92 (7.5%) hydrogel crosslinked with equimolar DTT and 0.5 mM DAS as a photoinitiator. Constructs were fabricated via two-photon polymerization (2PP) at 90 mW (left) and 100 mW (right), demonstrating compatibility for cell-laden bioprinting. **(D)** Schematic overview of RNA-seq sample comparisons across 2D cultures, 3D spheroids, and 3D constructs at multiple time points. **(E)** KEGG pathway enrichment analysis reveals further upregulation of the mevalonate pathway in encapsulated ES cultures compared to 2D (fgsea, FDR ≤ 0.005). Asterisks denote statistical significance based on adjusted p-values: *p* < 0.05 (\**), p < 0.01 (**), p < 0.001 (*\*\*\*)**. (F)** qRT-PCR data showing differences in gene expression between cells cultured in standard 2D conditions and the same cells cultured in 3D-constructs for 14 days. Each of the three 3D biological replicates combines 5 individual constructs. **(G)** ES PDX-derived cells (IC-pPDX-87) embedded in gelNB92, 7.5% hydrogel with equimolar concentrations of crosslinker dithiothreitol (DTT) and 0.1 mM Li-TPO-L remained viable and formed expanding spheroids over time, as confirmed by LIVE/DEAD staining (green: live cells, red: dead). Cells spontaneously assembled into spheroids within the hydrogel constructs, similar to their behavior in liquid culture. **(H)** Targeted dose–response analysis of doxorubicin in PDX-derived 3D-constructs. Three concentrations, selected from prior cell line spheroid screening, were tested in hydrogel-embedded constructs formed from dissociated IC-pPDX-87 cells. Cell viability was assessed after 72 h of treatment using luminescence assays. Each data point represents a single construct (one per well); three constructs were tested per concentration. This focused approach enabled detection of the drug’s effective range while minimizing use of limited patient-derived material.

Next, we assessed the molecular characteristics of the 3D constructs compared to spheroids using RNA-seq on day 14 (**Fig. 2D**). Similar to trends observed in spheroids, the mevalonate pathway was upregulated, with this effect being further enhanced under embedded (3D construct) conditions (fgsea^33^ FDR-adjusted P-value <= 0.005; **Fig. 2E**).

We validated the differential expression of selected EWS::FLI1 target genes and other functionally relevant genes in 3D constructs by qRT-PCR (**Fig. 2F**). Genes were selected based on their relevance to ES biology or their involvement in key processes such as proliferation, adhesion, and tumor progression. Notably, we detected upregulation of *AURKB*, a target gene of EWS::FLI1 highly expressed in ES tumors^48^ and considered a promising therapeutic target in ES^49^. Similarly, *IGF1*, a prominent EWS::FLI1 target gene known to be downregulated in 2D culture^50^ showed increased expression in 3D-constructs. E-cadherin (CDH1), an intercellular adhesion molecule crucial for cell adhesion and aggregation and metastasis^51^, was also elevated, consistent with enhanced cell-cell interactions in the 3D environment. In parallel, *TAZ/WWTR1* and downstream YAP/TAZ target genes (*SERPINE1*, *CYR61* and *CTGF)*, which regulate cellular plasticity^52^, proliferation, migration, and differentiation in both stem cells and cancer, showed reduced expression. These patterns suggest that 3D culture conditions modulate mechanotransduction and transcriptional programs, resulting in a gene expression profiles distinct from those in conventional 2D systems.

Given that limited cell availability is a common challenge when working with primary patient samples, we sought to model this constraint deliberately and test whether 3D encapsulated constructs could still support robust ex vivo culture and drug testing. To this end, tumor cells from an ES PDX model (IC-pPDX-87^43,53^) were dissociated and embedded, with standard liquid culture serving as a control. Within the hydrogel, patient-derived cells successfully formed spheroids with minimal or no cell death, as confirmed with LIVE/DEAD staining, further supporting the suitability of this model for long-term ex vivo studies. We observed a steady increase in cell numbers over time, demonstrating that the hydrogel matrix provided a supportive environment for cell growth (**Fig. 2G**).

We next applied a two-step strategy to assess drug sensitivity. Initial screening of ES cell line spheroids across multiple concentrations identified informative ranges, which were then tested in hydrogel-embedded PDX-derived cultures. Using doxorubicin as proof of principle, this approach successfully captured its effective range in 3D patient-derived constructs (**Fig. 2H**). By leveraging insights from spheroid testing, this workflow enables efficient drug evaluation while minimizing the use of scarce patient material, making complex 3D models a feasible and informative platform for translational studies.

### ES cells display synergistic vulnerability to mevalonate pathway and BCL-2 inhibition

Given the pronounced deregulation of the mevalonate pathway in ES spheroids (**Figs. 1D, S2, S4A and B**), we hypothesized that targeting this pathway with statins (**Fig. 3A**) could be a promising therapeutic approach. We selected pitavastatin (**Fig. 3B**), based on its favorable pharmacokinetic profile, including clinically achievable plasma concentrations, an extended half-life compared to other statins^42^, and a low risk of drug-drug interactions due to minimal metabolism by CYP450 enzymes^42,54^ and lack of P-glycoprotein involvement^55^.

**Figure 3.**
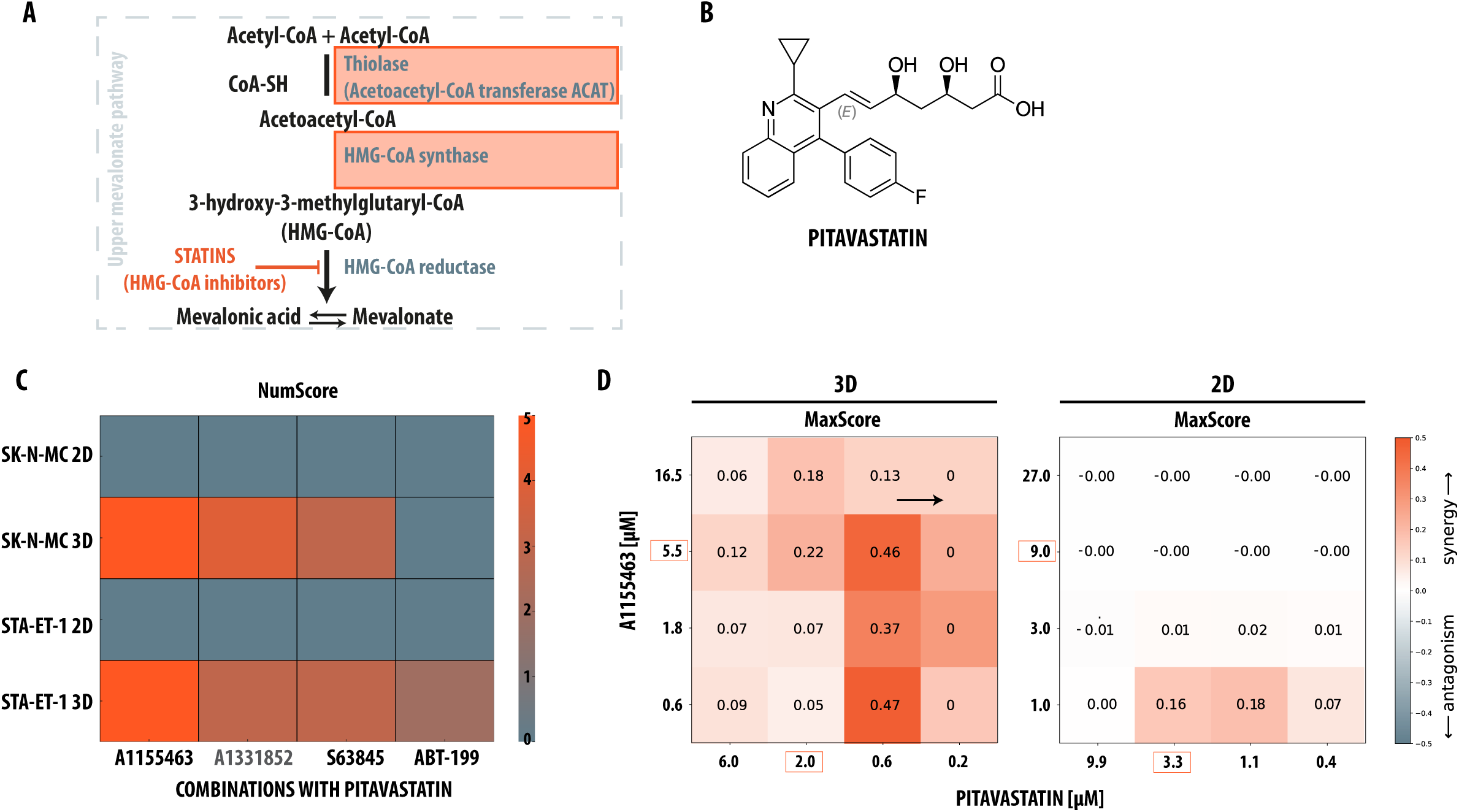
Synergistic interaction between pitavastatin and BCL-2 inhibitors in 3D Ewing sarcoma spheroid models. **(A)** Schematic overview of the upper mevalonate pathway illustrating statin-mediated inhibition of HMG-CoA reductase. **(B)** Chemical structure of pitavastatin, a potent HMG-CoA reductase inhibitor targeting mevalonate pathway. **(C)** Heatmap summarizing drug synergy in 3D Ewing sarcoma cultures. Combinations of pitavastatin with the BCL-2 inhibitor ABT-199 (venetoclax), BCL-xL inhibitor A1155463, or MCL-1 inhibitor S63845 were additive in 2D cultures (NumScore = 0) but showed marked synergy in 3D spheroids across multiple concentration points. NumScore indicates the number of wells in the dose matrix where the observed effect exceeds Bliss-predicted additivity. **(D)** Heatmap of MaxScore values for the SK-N-MC cell line, comparing 3D and 2D cultures. Left: High MaxScores in 3D cultures indicate strong synergy across concentration points. Right: Low MaxScores in 2D cultures highlight reduced synergistic interaction. Orange boxes denote the IC₅₀ concentrations of both pitavastatin and A1155463.

Based on our RNA-seq data and previous studies^56–60^, we next selected BCL-2 family protein inhibitors – small molecules that antagonize the pro-survival functions of anti-apoptotic proteins such as BCL2, Mcl1, and BCL-xL – for combination treatment with pitavastatin. To systematically assess drug interactions, we employed the Bliss independence model^61^, which assumes probabilistic independence between agents to quantify synergy. Although many mathematical descriptions of synergy exist^62^, we selected Bliss for its simplicity and numerical stability^63^. In our study, we calculated the deviation from Bliss independence across all data points within the factorial dose matrices used to evaluate pairwise combinations of pitavastatin with BH3 mimetics. This approach enabled us to capture the entire synergy landscape in each matrix. To facilitate interpretation and comparison across different drug combinations and cell lines, we derived two key summary metrics: (1) NumScore, representing the total number of concentration points exhibiting synergy (i.e. number of wells in the dose matrix that produced higher effect than the expected additive effect), and (2) MaxScore, indicating the maximum strength of synergy observed at any concentration point within the matrix. For NumScore, we set a synergy threshold at a Bliss excess score > 0.2, accounting for experimental noise^64^ and ensuring that the observed effects significantly exceed simple additivity.

Consistent with our hypothesis that combining pitavastatin with BCL-2 family inhibitors would produce enhanced anti-tumor effects specifically in the 3D context – where the mevalonate pathway is upregulated – all tested drug combinations (pitavastatin with the BCL-2 inhibitor ABT-199 [venetoclax], the BCL-xL inhibitor A-1155463, or the MCL-1 inhibitor S63845) showed only additive effects in 2D cultures, with NumScore frequently at zero, indicating no synergy at any concentration point. In contrast, 3D cultures exhibited significant synergy across multiple concentrations (**Fig. 3C**). To confirm that the observed synergy is specific to combined mevalonate pathway and BCL-xL inhibition, we tested a second BCL-xL inhibitor, A-1331852, and again observed a 3D-specific synergy in both cell lines (**Fig. 3C**). Although A-1331852 is more selective for BCL-xL^65^, its synergy was weaker than that of A-1155463, potentially reflecting broader target engagement by the latter.

As previously reported^59^, MCL1 and BCL-xL are key therapeutic targets in ES, with broad expression of *MCL1* and *BCL2L1* (encoding BCL-xL) across patient samples. This pattern was particularly pronounced in SK-N-MC cells, which displayed substantially higher NumScore and MaxScore values in 3D compared than the consistently low scores in 2D (**Fig. 3C,D**), highlighting the enhanced synergy observed in the 3D setting. Notably, ABT-199 did not exhibit synergy, despite reports of its synergistic effect with statins in leukemia^56^, likely reflecting the greater BCL-2 dependency in hematological malignancies compared with solid tumors^66^. Overall, the strongest and most consistent synergy was observed with A-1155463 across both ES cell lines.

To explore whether the observed synergy correlates with 3D-induced upregulation of the mevalonate pathway, we expanded our analysis to two OS cell lines (OS-143B and STA-OS-5). The steroid biosynthesis pathway – a key branch of the mevalonate pathway – undergoes more extensive alteration in ES spheroids than in OS, with distinct key components affected. OS cells exhibited minimal changes in steroid biosynthesis when transitioning from 2D to 3D, whereas ES cells underwent substantial alterations (**Fig. S4B**). Functionally, synergy between pitavastatin and BCL-xL inhibitor was weak or absent in STA-OS-5, often undetectable in some replicates or reflected by low NumScore (**Fig. S4C**). When synergy was observed, it paradoxically favored 2D over 3D conditions. In contrast, OS-143B exhibited modest synergy that was more consistent with the ES trend, showing slightly enhanced effects in 3D. Nevertheless, MaxScore comparisons revealed that ES cell lines consistently achieved higher peak synergy than OS models (**Fig. S4D**).

Together, these results suggest that while MaxScore captures peak synergy, differences in both MaxScores and NumScores underscore the differing capacities of ES and OS cells to achieve synergistic effects. The robust and 3D-specific synergy observed in ES appears closely tied to mevalonate pathway upregulation – a metabolic adaptation that is less prominent or differently regulated in OS. The pronounced shift in ES between 2D and 3D further reinforces the role of microenvironmental context in revealing metabolic vulnerabilities.

## Discussion

In this study, we present a robust 3D culture platform for focused drug screening in pediatric bone sarcomas, with a comparative focus on ES and OS. By selecting cell lines capable of consistent, single spheroid formation under low-attachment conditions and optimizing drug administration workflows, we developed a reproducible and scalable system suitable for automation, which can inform drug testing in more complex 3D models, including matrix-embedded and bioprinted constructs.

A key strength of our platform is its reproducibility and capacity to generate high-resolution dose-response data, enabling precise quantification of drug potency and efficacy. While 3D cultures introduce additional complexity, cost, limited throughput, and interpretation challenges, they more faithfully recapitulate in vivo tumor features, including nutrient gradients, hypoxia, and altered metabolism^67^. Consistent with this, ES spheroids displayed upregulation of the mevalonate pathway, a metabolic adaptation commonly observed in cancers^68^ and previously reported in 3D colon cancer models^69^. Morphological analyses revealed phenomena such as spheroid expansion at high doxorubicin concentrations, likely reflecting oncotic swelling and necrotic core expansion^32^, highlighting the value of integrating phenotypic and viability readouts.

Using this platform, we identified a potent and selective synergy between pitavastatin, a mevalonate pathway inhibitor, and BCL-xL inhibitors in ES spheroids, detectable only under 3D conditions and absent in conventional 2D assays. Synergy was initially assessed via established Bliss independence model, and to facilitate interpretation, we developed two complementary metrics, NumScore and MaxScore, summarizing the overall extent and peak magnitude of synergy across concentration ranges. Transcriptomic analyses revealed that steroid biosynthesis, a key branch of the mevalonate pathway, was markedly upregulated in ES under 3D conditions, whereas OS engaged alternative metabolic programs, highlighting distinct adaptive strategies and reinforcing the importance of physiological context in uncovering metabolic vulnerabilities. Although BCL-xL inhibition is clinically limited by thrombocytopenia^70^, emerging strategies such as the PROTAC DT2216^71^ that selectively degrades BCL-xL in a manner, may enable translational application in combination with pitavastatin.

Despite their advantages, 3D models remain simplifications of the in vivo tumor environment, lacking vasculature, mechanical forces, and dynamic fluid flow^72^. To accommodate the diverse growth behaviors of bone sarcomas, we implemented two complementary approaches – scaffold-free spheroids and matrix-embedded constructs – facilitating spatial organization, improved viability of primary cells and compatibility with patient-derived materials.

Overall, our study demonstrates that physiologically relevant 3D culture systems can uncover clinically relevant drug interactions masked in conventional 2D assays. The identified 3D-specific synergy between pitavastatin and BCL-xL inhibitors in ES highlights the importance of metabolic context and supports further development of targeted therapies for pediatric bone sarcomas.

## Materials and Methods

### 2D cell culture

ES cell lines (TC-71, A673, STA-ET-1, SK-N-MC) and osteosarcoma (OS) cell lines (STA-OS-1, STA-OS-2, STA-OS-3, STA-OS-5, U2OS, OS143B) were cultured in RPMI medium (RPMI Medium 1640 (1x) + GlutaMax, Gibco, Cat. #61870-010) supplemented with 10% fetal calf serum (FCS, Gibco, Cat. #A5256701) and 1% penicillin-streptomycin (PAN Biotech, Cat. #P06-07100) at 37°C in a 5% CO₂ atmosphere. For standard 2D culture, STA-ET-1 cells require fibronectin-coated surfaces (Fibronectin (pure), Roche, Cat. #11080938001). SK-N-MC cell line authentication was conducted via short tandem repeat (STR) profiling at Microsynth. Routine mycoplasma testing was carried out using the MycoAlert Mycoplasma Detection Kit (Lonza, Cat. #LT07-318).

In our study, we mostly focused on the SK-N-MC and STA-ET-1 cell lines, both detailed in Ottaviano et al.^73^. The SK-N-MC line, derived from an Askin tumor in the supra-orbital region, displays the *EWSR1::FLI1* gene fusion^3,74,75^ and a *TP53* c.170_572del mutation^73^. Despite lacking functional p53, it can activate p21 WAF1 expression^76^. The STA-ET-1 line exhibits the *EWSR::FLI1* fusion^77^, a homozygous *CDKN2A* deletion^1^, and no detected *TP53* mutations.

### Spheroid cell culture

Prior to spheroid initiation, cells were passaged at least twice and no more than ten times to minimize culture-induced variability. For spheroid assays, cells were seeded into 96-well plates coated with 1.5% agarose prepared from LE Agarose (Biozym, Germany; Cat. #840004), which we found to be critical for consistent spheroid formation. Agarose was prepared by gradual heating in RPMI Medium 1640 (1x) + GlutaMax (Gibco, USA; Cat. #61870-010) supplemented with 1% penicillin-streptomycin (PAN Biotech, Germany; Cat. #P06-07100). Notably, agarose was melted in serum-free medium with antibiotics, avoiding fetal calf serum during spheroid embedding. Unlike prior protocols requiring autoclaving, agarose preparation did not involve this step, which did not compromise sterility or consistency. Attempts to use alternative agarose products were unsuccessful, making this specific formulation essential for reproducible spheroid formation.

Cells were seeded at densities ranging from 1×10³ to 1×10⁴ cells per well in a final volume of 100 µL RPMI medium (RPMI Medium 1640 (1x) + GlutaMax, Gibco, Cat. #61870-010) supplemented with 10% fetal calf serum (FCS, Gibco, Cat. #A5256701) and 1% penicillin-streptomycin (PAN Biotech, Cat. #P06-07100). Attempts to reduce the seeding volume impaired spheroid formation, highlighting the importance of adequate medium volume for aggregation. Plates were incubated at 37°C in 5% CO₂ for 72 hours, resulting in a single spheroid per well of approximately 400 µm in diameter. Media was exchanged by gently replacing half the volume with fresh medium to minimize spheroid disturbance.

### Spheroid staining and cryosectioning

Spheroid viability was assessed using the LIVE/DEAD® Viability/Cytotoxicity Kit (Invitrogen, Thermo Fisher Scientific, USA; Cat. #L3224). Spheroids were incubated with 4 µM EthD-1 and 0.625–1.25 µM Calcein-AM (optimized for cell line and spheroid density) in 200 µL per well (96-well plate) and incubated for 30–90 minutes at 37 °C. Following incubation, spheroids were gently washed three times with PBS (1X) (Gibco, Thermo Fisher Scientific, USA; Cat. #14190144) to remove excess dye. Fluorescence imaging was performed using SP8 X WLL confocal microscope system (Leica, Germany) with separate HyD detectors and excitation/emission settings of 494/517 nm for Calcein-AM (live, green) and 528/617 nm for EthD-1 (dead, red).

For proliferation analysis, spheroids were cryosectioned and stained with a primary antibody against Ki67 (Abcam, Cambridge, UK; Cat. #16667, 1:1000 dilution). Sections were washed and mounted using VECTASHIELD® Antifade Mounting Medium containing DAPI (Vector Laboratories, Newark, CA, USA, Cat. #H-1200-10), to counterstain nuclei (blue) and preserve fluorescence. Ki67 expression and nuclear localization were imaged on the same confocal system.

For drug penetration studies, spheroids treated with doxorubicin were imaged on day 7 without additional staining, using doxorubicin’s intrinsic autofluorescence (excitation 470 nm, emission 595 nm) on the SP8 X WLL confocal microscope (Leica, Germany).

### Liquid culture – PDX-derived cells

IC-pPDX-87 was generated at Institut Curie from patients under an Institutional-Review-Board-approved protocol (OBS170323CPP ref3272; dossier No. 2015-A00464-45)^43^. Tumor fragments from IC-pPDX-87 were dissociated following the protocol described by Stewart et al^78^. The resulting IC-pPDX-87 patient-derived xenograft (PDX) cells were cultured in DMEM/F-12 medium with GlutaMAX™ (Gibco, Thermo Fisher Scientific, USA, Cat. #11514436), supplemented with 1% B-27 (50X, Gibco, Thermo Fisher Scientific, USA, Cat. #11530536) and 1% Penicillin-Streptomycin (PAN Biotech, Germany, Cat. #P06-07100). Cells were maintained at 37 °C in a humidified incubator with 5% CO₂ and spontaneously formed spheroids. For passaging, cells were dissociated using Accutase (Pan Biotech, Germany, Cat. #P10-21100).

### Cell-embedded constructs

We optimized conditions for the effective cultivation of ES cells and ES PDX-derived cells within a gelatin-norbornene (gel-NB^79^) hydrogel matrix. The optimal formulation consisted of 7.5% gel-NB with a high degree of substitution^80^ (92%), 0.1 mM Li-TPO-L photoinitiator and dithiothreitol (DTT) as a crosslinker at an equimolar thiol-ene ratio. Under these conditions, cells remained viable and proliferative for 14–21 days, with medium replaced every 2–3 days.

Approximately 9 × 10^6^ PDX-derived tumor cells as a single suspension were used to generate 20 hydrogel constructs (∼5 × 10^5^ cells per construct). A 30 µL aliquot of the solution was pipetted onto a methacrylated^81^ glass bottom dish (ibidi, Germany) and placed in the UV curing chamber (Emission dose of 1 J over 4.5 min, 365 nm, UV Crosslinker AH, Boekel Scientific). After crosslinking, constructs were washed twice with medium to remove any residual material and cultured at 37°C with 5% CO_2_ in a humid atmosphere.

Cell embedding using 2-photon-polimerisation (2PP) was performed as described previously^46,82^. Gel-NB was prepared as above, except that 0.5 mM of the DAS^47^ served as the photoinitiator. Next, 30 µL of the obtained solution was pipetted into a silicone mold (6 mm diameter, 1 mm height) placed on the methacrylated glass bottom dish. The resulting constructs were fabricated as 50 µm-pore beehive structure (400 x 400 µm^2^), using a 10x/0.3 Plan-Apochromat objective (Zeiss, Oberkochen, Germany). After crosslinking, the constructs were processed as described above.

Cell viability was assessed using the LIVE/DEAD™ Viability/Cytotoxicity Kit (Invitrogen, Thermo Fisher Scientific, USA), following in-house adapted protocol^44^. After media removal, both construct and cell pellets were rinsed three times with sterile phosphate-buffered saline (PBS; Sigma-Aldrich, USA). A staining solution containing 0.2 μM Calcein AM (live cell marker) and 0.6 μM propidium iodide (dead cell marker) was applied, followed by a 20-minute incubation at 37°C. Post-incubation, samples were washed three times with PBS and transferred to 35 mm glass-bottom dishes (ibidi, Germany) for imaging using a laser scanning microscope (LSM 700, Zeiss, Germany). Fluorescence was detected using an excitation/emission filter of 488/530 nm for live cells (green) and 530/580 nm for dead cells (red).

### Compounds

S63845 and A-1155463 were obtained from Selleck Chemicals (Houston, TX, USA) and TargetMol (Boston, MA, USA). ABT-199 (venetoclax) and A-1331852 were purchased from THP Medical Products (Vienna, Austria) and TargetMol (Boston, MA, USA). Simvastatin was acquired from TargetMol (Boston, MA, USA), while pitavastatin was obtained from Cayman Chemical (Ann Arbor, MI, USA). Doxorubicin, etoposide, and vincristine were purchased from Selleck Chemicals (Houston, TX, USA).

### Viability assays

Cell viability was assessed using the CellTiter-Glo® Luminescent Cell Viability Assay (Promega, Madison, WI, USA, Cat. #G7570) or the CellTiter-Glo® 3D Cell Viability Assay (Promega, Madison, WI, USA, Cat. #G9681), according to the manufacturer’s instructions. One day prior to drug treatment, cells were seeded into 96-well viewplates (Revvity, Inc., Waltham, MA, USA, Cat. #6005181) at densities optimized for each cell line, typically ranging from 1×10³ to 1×10⁴ cells per well in 50 µL of medium, and allowed to adhere overnight. For STA-ET-1 cells, fibronectin-coated plates (Fibronectin (pure), Roche, Basel, Switzerland, Cat. #11080938001) were used for seeding.

For 3D cultures, cells were passaged twice before seeding into 1.5% agarose-coated white 96-well viewplates (Revvity, Waltham, MA, USA, Cat. #6005181) at 1×10³ to 1×10⁴ cells per well in 100 µL of medium. These were incubated for 72 hours to allow formation of robust spheroids. After spheroid formation, 50 µL of excess medium was carefully removed from each well using a multichannel pipette.

Compounds were serially diluted in standard 96-well plates and subsequently transferred to the cell plates to generate a two-drug matrix (final volume: 50 µL per well). Plates were incubated at 37°C with 5% CO₂ for an additional 72 hours. To assess viability, CellTiter-Glo® (diluted 1:4 in 1× DPBS) or CellTiter-Glo® 3D (for spheroids) was added to each well (100 µL), followed by incubation on an orbital shaker for 10 minutes for 2D cultures and 30 minutes for 3D culture, followed by incubation in the dark (15 minutes for 2D; 30 minutes for 3D). Luminescence was measured using either an EnSpire plate reader (PerkinElmer, Waltham, MA, USA) or a Spark Cyto plate reader (Tecan Group, Männedorf, Switzerland). The final DMSO concentration in assay wells was kept below 0.2%. All conditions were tested in triplicate. For automated imaging, Operetta CLS high-content imager (PerkinElmer now Revvity, Waltham, MA, USA) was used, with the following settings: 5x air objective and brightfield imaging (40 ms exposure, 10% intensity, 4 planes of 250µM to cover various suspense levels). For determining sphere size in the Harmony Software version 4.9 (PerkinElmer now Revvity, Waltham, MA, USA) images were pre-filtered by the modification of inversion to enable spheroid detection by intensity threshold. Size dimensions of spheroid were approximated through the projection of the spheroid along the z-axis onto the x-y-plane, the footprint area in µm².

### Drug synergy analysis

To systematically assess drug interactions, we employed the Bliss independence model^61^, which assumes that drugs act independently when the combined effect equals the product of their individual effects. This model was selected for its simplicity, probabilistic foundation, and numerical stability^63^. For each drug, 72-hour IC₅₀ values were first determined for each cell line, and dilution series were designed using the IC₅₀ and Hill slope to create symmetric dose-response matrices. Typically, a 3-fold dilution was applied, centered around the IC₅₀, starting from 3× or 9× IC₅₀. Each drug pair was tested using triplicate matrices in separate 96-well plates (ViewPlate-96, Revvity, Waltham, MA, USA, Cat. #6005181). Cell viability was measured using either the CellTiter-Glo® Luminescent Cell Viability Assay (Promega, Cat# G7570) or CellTiter-Glo® 3D Assay (Promega, Cat# G9681). Bliss-predicted effects were computed for each concentration pair and compared to observed viability to classify interactions as additive (Bliss score = 0), synergistic (>0), or antagonistic (<0). To summarize and compare combinations across drugs and cell lines, we extracted two metrics: (i) *NumScore* – the number of concentration points showing synergy, and (ii) *MaxScore* – the highest observed Bliss excess in a matrix. A Bliss threshold >0.2 was applied to define synergy, accounting for experimental noise^64^ and ensuring robustness of synergy detection.

### RNA-seq analysis

For all comparisons, 2D and 3D cultures were grown in identical medium to control for effects of culture conditions on gene expression. To prepare samples for RNA sequencing, total RNA was isolated using RNeasy Mini Kit (#74106, QIAGEN) or RNeasy Plus Universal Mini Kit ((#73404, QIAGEN) for embedded constructs, following manufacturers’ protocols. For embedded samples, three cell-loaded constructs were pooled per condition to ensure sufficient RNA yield. Constructs were shock frozen in liquid nitrogen and grinded directly in microcentrifuge tubes using a micro-pestle. Subsequently, 900 µL of QIAzol Lysis Reagent (QIAGEN) were added to the tube and RNA was purified using the RNeasy Plus Universal Mini Kit according to the manufacturer’s instructions. RNA quantity and quality were assessed prior to library preparation. Sequencing was performed in 50 bp single-end read mode on an Illumina HiSeq instrument at the Biomedical Sequencing Facility (BSF) at the CeMM Research Center for Molecular Medicine of the Austrian Academy of Sciences.

Raw sequencing data were processed using the quantseq and rnaseq pipelines in looper v0.12.3 and pypiper v0.10.0^83^ Briefly, reads were trimmed off adapter using bbduk/bbmap v38.87 (parameters: *k=13 ktrim=r useshortkmers=t mink=5 qtrim=r trimq=10 minlength=20*; for Quant-seq) or trimmomatic v0.36^84^ (parameters: ILLUMINACLIP:adapterfile:2:10:4:1:true SLIDINGWINDOW:4:1 MAXINFO:16:0.40 MINLEN:30; for RNA-seq), aligned against the XXX reference genome using STAR v2.7.3a^85^ (parameters: *--outFilterType BySJout --outFilterMultimapNmax 20 --outSAMunmapped Within --alignIntronMax 1000000 --alignMatesGapMax 1000000 --alignIntronMin 20 --alignSJDBoverhangMin 1 --alignSJoverhangMin 8 --outFilterMismatchNmax 999* [common]; *--outFilterMismatchNoverLmax 0.1 --outSAMmapqUnique 60* [only Quant-seq]; *--sjdbScore 1 --outFilterMismatchNoverLmax 0.04* [only RNA-seq]), and subsequently loaded into R v4.1.3 using Rsubread::featureCounts v2.8.2^86^ for all downstream data processing. In doing so, alignments to the mitochondrial genome and sex chromosomes were removed. The samples “BRS_3_3_2d" and "BRS_4_4_2d” were removed because they came from a separate experimental run. Only genes detected in at least four samples with at least five reads were considered for further analysis. Differential expression analysis was performed using DESeq2 v1.34.0^87^ (parameters: *sfType=”poscounts”, betaPrior=FALSE*). Genes with an FDR-adjusted P-value less or equal to 0.05 and an absolute log_2_ fold change greater or equal to 0 were considered differentially expressed. For pathway enrichment analysis, we used fgsea v1.20.0^88^ the reversed log_2_ fold change as a ranking criterion (parameters: *min_size=10, max_size=1000*), and the KEGG and MatrisomeDB databases. Pathways with an adjusted P-values less or equal to 0.005 in comparison between ES spheroids at days 4 or 7 vs. 2D were selected for display in all plots. Plots were generated using standard R functions, ggplot2 v3.3.5, pheatmap v1.0.12, and ComplexHeatmap v2.10.0.

## Acknowledgements

We gratefully acknowledge financial support from the Austrian Science Fund (FWF) and the Federal Ministry of Education, Science and Research (BMBWF) under FWF Project No. P 35353 (H.K.). This work was also supported through: Austrian Research Promotion Agency (FFG) project 7940628 (Danio4Can) (M.D.). We thank Prof. Sandra Van Vlierberghe (Polymer Chemistry and Biomaterials Group, Centre of Macromolecular Chemistry, Department of Organic and Macromolecular Chemistry, Ghent University, Ghent, Belgium) for kindly providing Gel-NB. We also thank Eva Scheuringer for assistance with Operetta handling and image analysis, and Agnes Dobos for support with two-photon polymerization (2PP) printing.

---

**Supplementary Figure S1.**
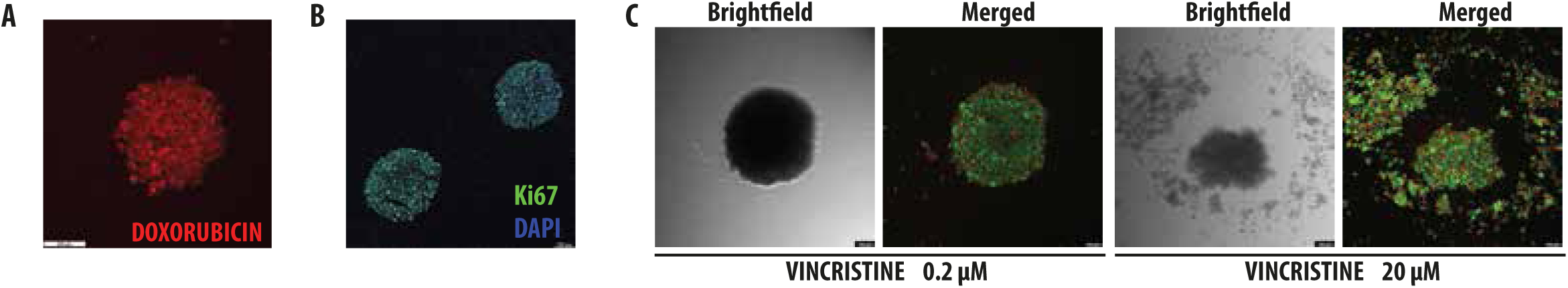
Visualization of proliferation, viability, and drug penetration in sarcoma models using confocal microscopy. **(A)** Representative central optical section (middle slice) of a confocal Z-stack of doxorubicin-treated STA-OS-5 spheroids on day 6, demonstrating drug penetration via intrinsic autofluorescence of doxorubicin. **(B)** Cryosections of OS143B spheroids at day 4, stained for DAPI (nuclei, blue) and Ki67 (proliferation marker, green). **(C)** LIVE/DEAD viability assay of vincristine-treated STA-ET-1 cells showed no effect at 0.2 µM, while 20 µM induced substantial cell disruption. Live cells fluoresced green, dead cells red.

**Supplementary Figure S2.**
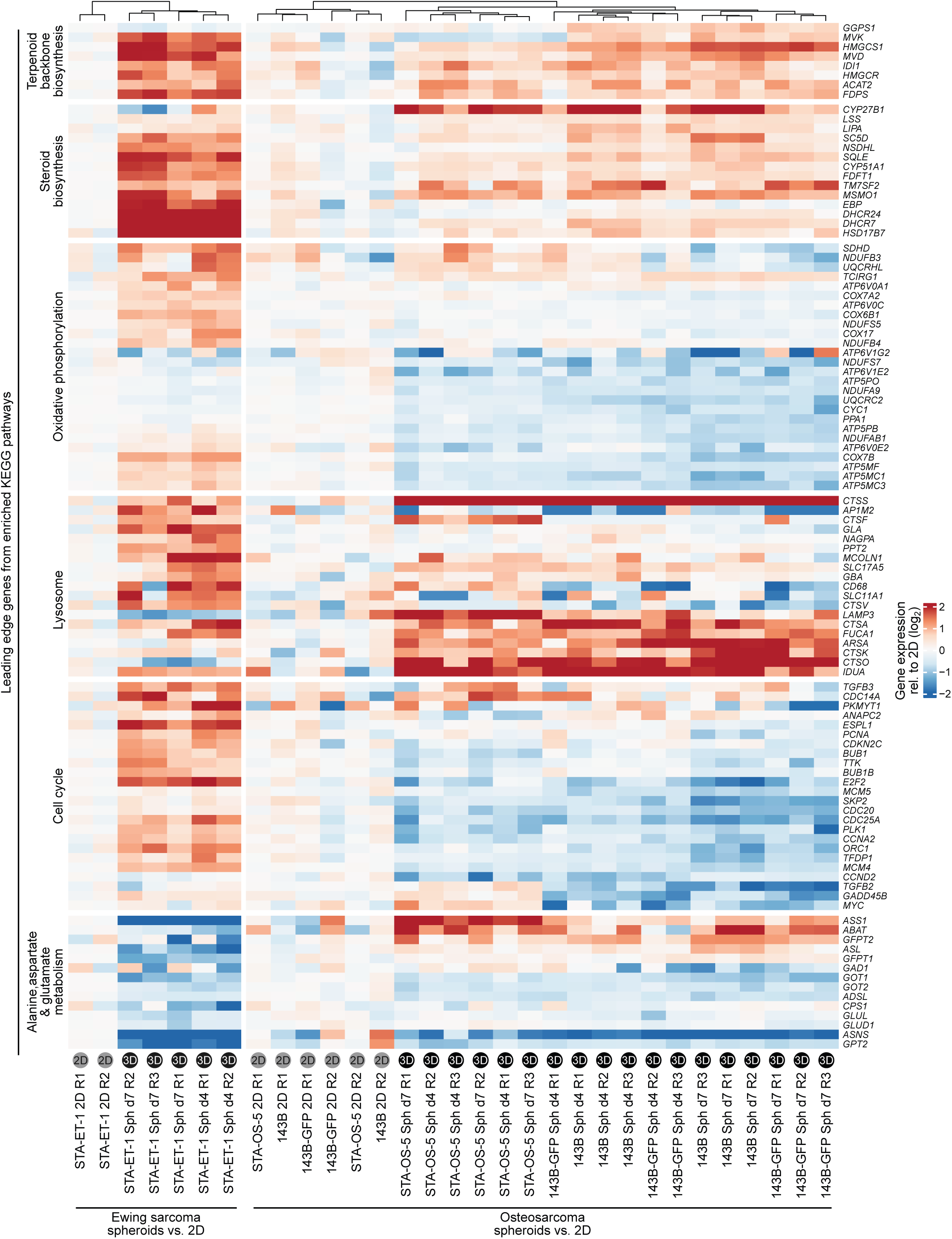
Pathway-level transcriptomic shifts in 3D ES and OS spheroids. Gene set enrichment analysis (GSEA) of RNA-seq data comparing 3D spheroids (d4 and d7) versus 2D cultures in Ewing sarcoma and osteosarcoma cell lines. Significant enrichment was observed for lysosomal activity, steroid biosynthesis, terpenoid backbone biosynthesis (components of the mevalonate pathway), and other metabolic and stress-response pathways. Enrichment was calculated using fgsea^33^ with FDR-adjusted p-values ≤ 0.005.

**Supplementary Figure S3.**
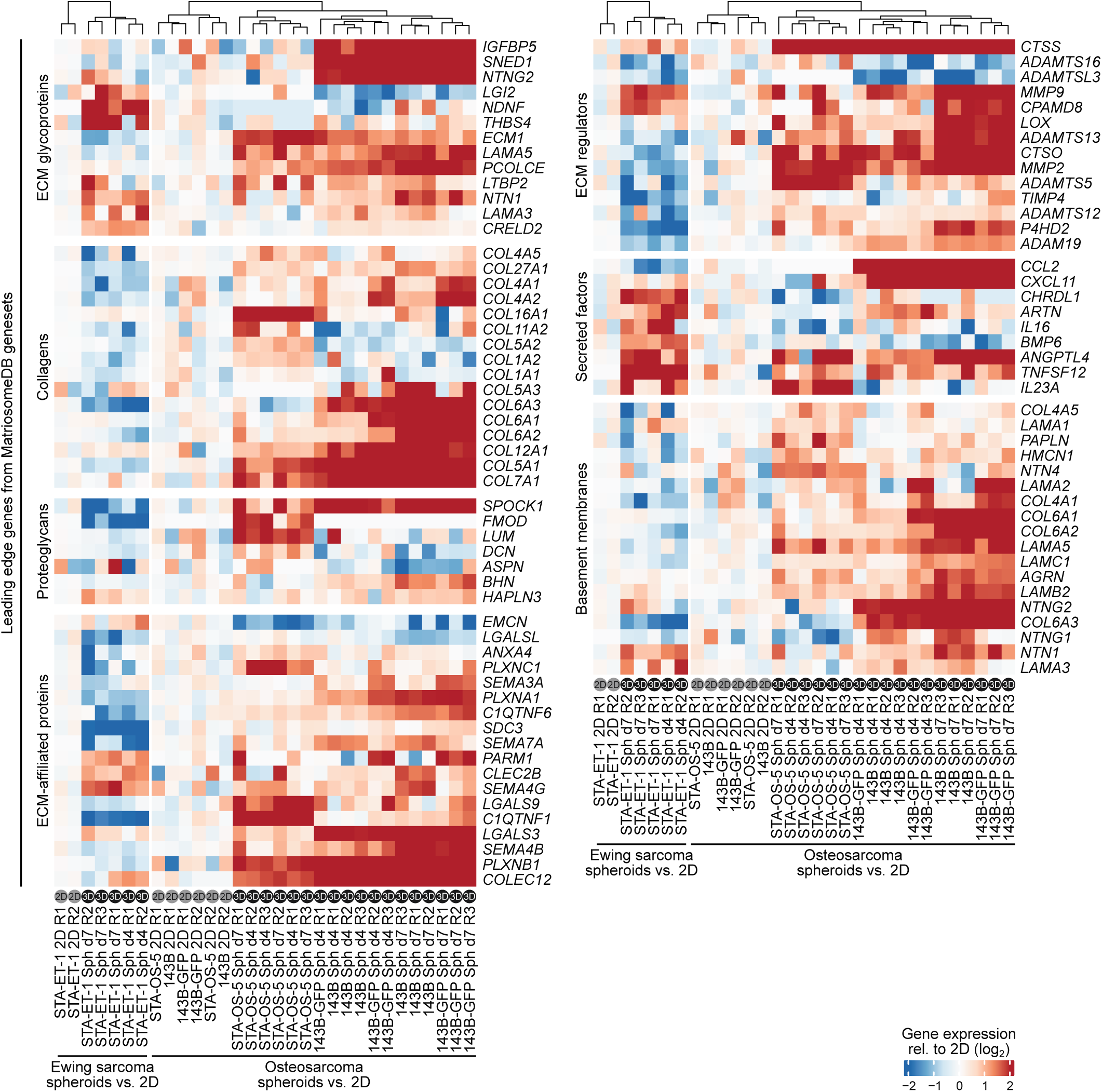
Differential expression of matrisome-associated genes and secreted factors in 3D cultures. Heatmap showing differential expression of matrisome genes and secreted factors annotated in MatrisomeDB^38^. Osteosarcoma spheroids showed broad upregulation of genes encoding extracellular matrix (ECM) components and regulators. In contrast, Ewing sarcoma spheroids showed more selective upregulation of ECM glycoproteins and secreted factors such as IL23A, BMP6, and TNFSF12 when compared to 2D cultures.

**Supplementary Figure S4.**
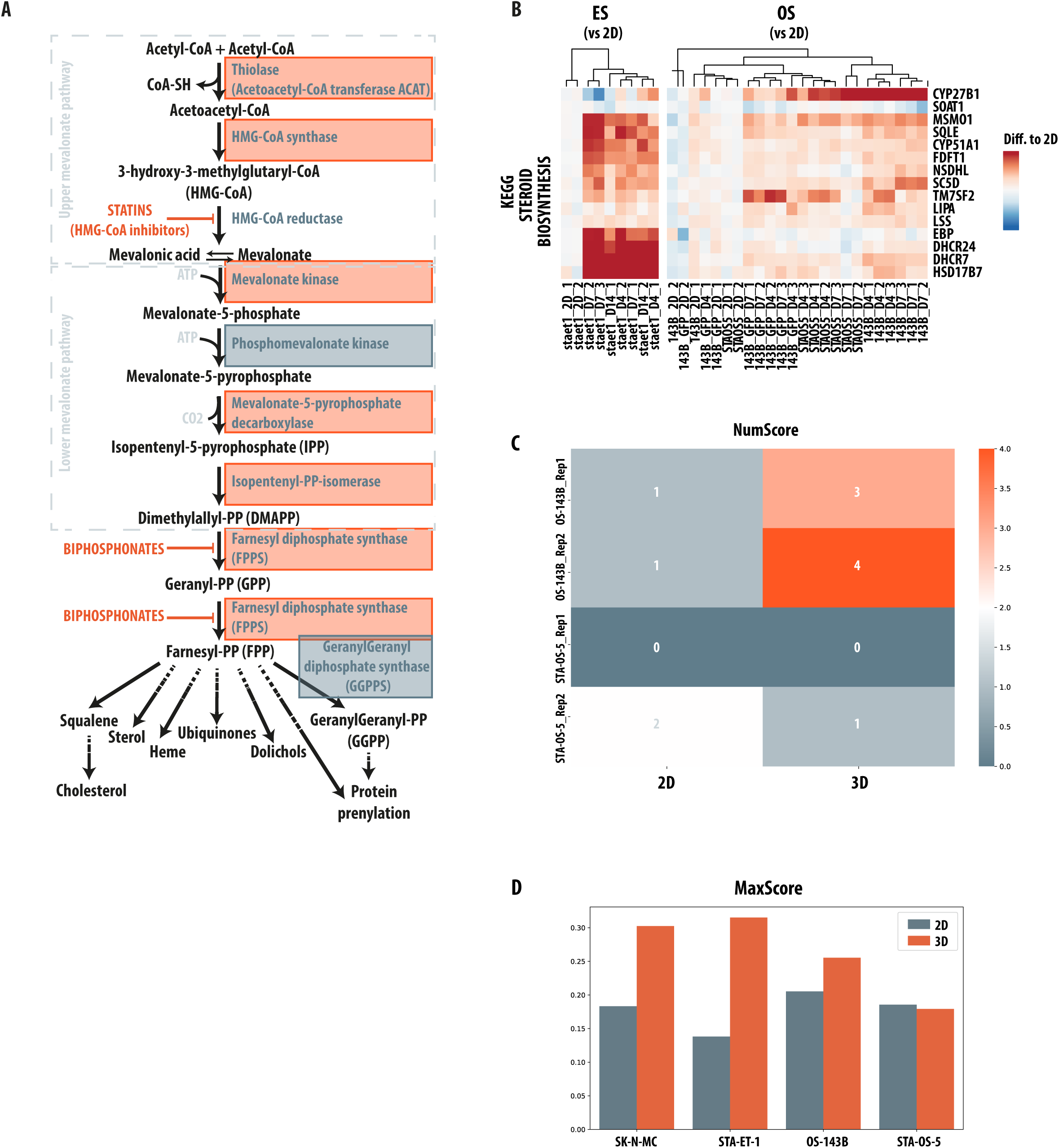
Differential regulation of the mevalonate pathway and drug synergy in ES and OS 3D cultures. **(A)** Schematic of the mevalonate pathway showing differential gene expression in ES spheroids versus 2D cultures. Upregulated enzymes are marked in red and downregulated enzymes in blue, highlighting widespread pathway activation, particularly within nodes involved in steroid biosynthesis and isoprenoid production. **(B)** Heatmaps comparing gene expression changes in the KEGG steroid biosynthesis pathway between 2D and 3D cultures for OS and ES cell lines. OS models showed limited and variable deregulation, whereas ES spheroids displayed consistent upregulation of key components, supporting enhanced pathway activity in 3D. Color scale indicates normalized enrichment score (NES), which reflects the degree of enrichment of a gene set normalized for gene set size and multiple testing. **(C)** Heatmap summarizing synergy (via NumScore) of pitavastatin and A1155463 in 2D vs. 3D osteosarcoma cultures. Compared to ES, synergy was weaker or absent in osteosarcoma, with STA-OS-5 showing a reversed effect, where activity was stronger in 2D cultures. **(D)** Comparison of MaxScore values between 3D and 2D cultures in two ES and two OS cell lines. Overall, MaxScore was higher in ES, with more pronounced 2D vs. 3D differences. STA-OS-5 exhibited a mild reversed effect, with stronger activity in 2D.

## Notes

### Competing Interest Statement

The authors have declared no competing interest.

